# Transmission Dynamics of Human Herpesviruses and Other Blood DNA Viruses from Whole Genome Sequences of Families

**DOI:** 10.1101/2022.01.31.478555

**Authors:** Brianna Chrisman, Chloe He, Jae-Yoon Jung, Nate Stockham, Kelley Paskov, Peter Washington, Dennis P. Wall

## Abstract

While hundreds of thousands of human whole genome sequences (WGS) have been collected in the effort to better understand genetic determinants of disease, these whole genome sequences have rarely been used to study another major determinant of human health: the human virome. Using the unmapped reads from WGS of 1,000 families, we present insights into the human blood DNA virome. In addition to extensively cataloguing the viruses detected in WGS of human whole blood and lymphoblastoid cell lines, we use the family structure of our dataset to show that household drives transmission of many microbes. We also identify several cases of inherited chromosomally integrated herpes 6A and 6B and locate candidate integration sequences for these cases. We document genetic diversity within exogenous and integrated HHV species and within integration sites of HHV-6. Finally, in the first observation of its kind, we present evidence that suggests widespread *de novo* HHV-6B integration and HHV-7 episome replication in lymphoblastoid cell lines. These findings show that the unmapped read space of WGS may be a promising avenue for virology research.

## 2 Background

### 2.1 A Wealth of Whole Genome Sequencing Data

As the cost and speed of whole genome sequencing (WGS) continues to improve, many research institutions have undertaken large scale whole genome sequencing studies in an effort to better understand genetic determinants of human diseases [51, 33, 27, 11]. While high coverage (>30x) WGS produces several hundred gigabytes of raw data per sample [36], in many pipelines up to 30% of these reads go unused because they fail to align to the human reference genome. [45]. These unmapped reads may originate from non-reference human DNA sequences, organic reagents and contamination, and human viruses.

### 2.2 Applications of the Blood Virome

Meanwhile, the last decade of advances in sequencing has also empowered the field of metagenomics and the study of the human microbiome, areas where next-gen sequencing technologies have allowed for the rapid characterization of bacteria, small eukaryotes, and viruses that inhabit human environments. While much of microbiome and virome research has focused on the gut microbiome [50, 32, 20] which has clear communication links between the human digestive system, nervous system and immune system, recently it has been suggested that microbiota with low microbial loads may also play novel roles in disease [12, 5, 54]. One such microbiota is human blood, and it is still up for debate whether or not there is a healthy blood microbiota or whether the presence of bacteria inherently indicates disease. [6, 46, 22]. Several studies have investigated the blood bacteriome in an attempt to understand the healthy blood bacterial microbiome, but only a handful studies have attempted to characterize the human blood virome and have been done in mostly diseased cohorts [37, 10, 4]. Furthermore, despite the importance of the blood virome in blood transfusion and stem cell transplant safety [18, 52], research in emerging pathogens [25, 15], and immune system regulation [16], to our knowledge only one study has analyzed the blood virome on the scale of thousands of individuals [35].

### 2.3 Intrafamilial Transmission and Inheritance of Viruses

The early stages of the SARS-CoV-2 pandemic underscored how important it is to understand transmission patterns for different viruses. [31]. Particularly in the case of intrafamilial transmission, non-sexual transmission between members of the same household, a better understanding of such might help families take steps to mitigate the risk of infection. Several known bloodborne viruses with high disease risk, such as betapapillomaviruses and hepatitis C [53, 40], show evidence of intrafamilial transmission. Furthermore, some human herpesviruses (HHV) have the ability to integrate into host genomes, and ancient integration events of human herpesvirus 6A and human herpesvirus 6B have persisted as a relatively common genotype, displaying common inheritance patterns. The prevalence and integration patterns of herpes 6A and 6B are not yet fully understood, though inherited chromosomally integrated herpesviruses (iciHHV) may place a role in cardiovascular disease [29, 19].

The iHART dataset [44] contains whole genome sequences from whole blood (WB) or lymphoblastoid cell lines (LCLs) from over 4,500 individuals from 1,000 different nuclear families with multiplex autism. Originally curated to understand the genetic determinants of autism, the iHART dataset has become valuable not only for autism research, but because its unique family structure allows for understanding of inheritance patterns that cannot be done with case-control cohorts [42, 7, 8]. In this study, we utilize the the family structure to better understand intra-family viral transmission and integration patterns. We characterize the human blood DNA virome, focusing particularly on intra-family transmission patterns and chromosomal integration and inheritance of herpesviruses.

## 3 Results

### 3.1 Unmapped Read Space Characterizes Prevalence and Abundance of Viruses

Using unmapped or poorly aligned reads from WGS of 4,569 individuals, we were able to reclassify reads to over 100 species of viruses. We show the top 50 most abundant viruses in Fig. 2, clustered by Spearman association across samples. Of note, we see four important categories of viruses: Human herpesviruses (HHV) 6A, 6B, and 7 are common blood viruses that are normally acquired during childhood. HHV-6 has the ability to integrate into host cells, and be inherited through ancient integration events in germline cells that are passed down mendellianly. HHV-6A, 6B, and 7 viral reads are likely true HHVs present in the blood, and we discuss the viral load profiles of HHV-6 and HHV-7 in depth later on. Lambda phage PhiX is a common reagent used in sequencing pipelines to calibrate Illumina machines and balance GC content. Reads classified as PhiX relatives are probably either mismappings to homologous regions, or contamination of the commercial PhiX reagents [35]. Similarly, Epstein Barr virus (EBV) or human gammaherpesvirus 4, is used to immortalize lymphoblastoid cell lines (LCLs), and so EBV and relatives are probably artifacts from the LCL immortalization pipeline. Torque Teno Viruses (TTV) and Erythroviruses are fairly common blood viruses that are usually acquired during childhood. We suspect the TTV and erythrovirus reads are probably true reads originating from an active TTV or erythrovirus infection.

**Figure 1:**
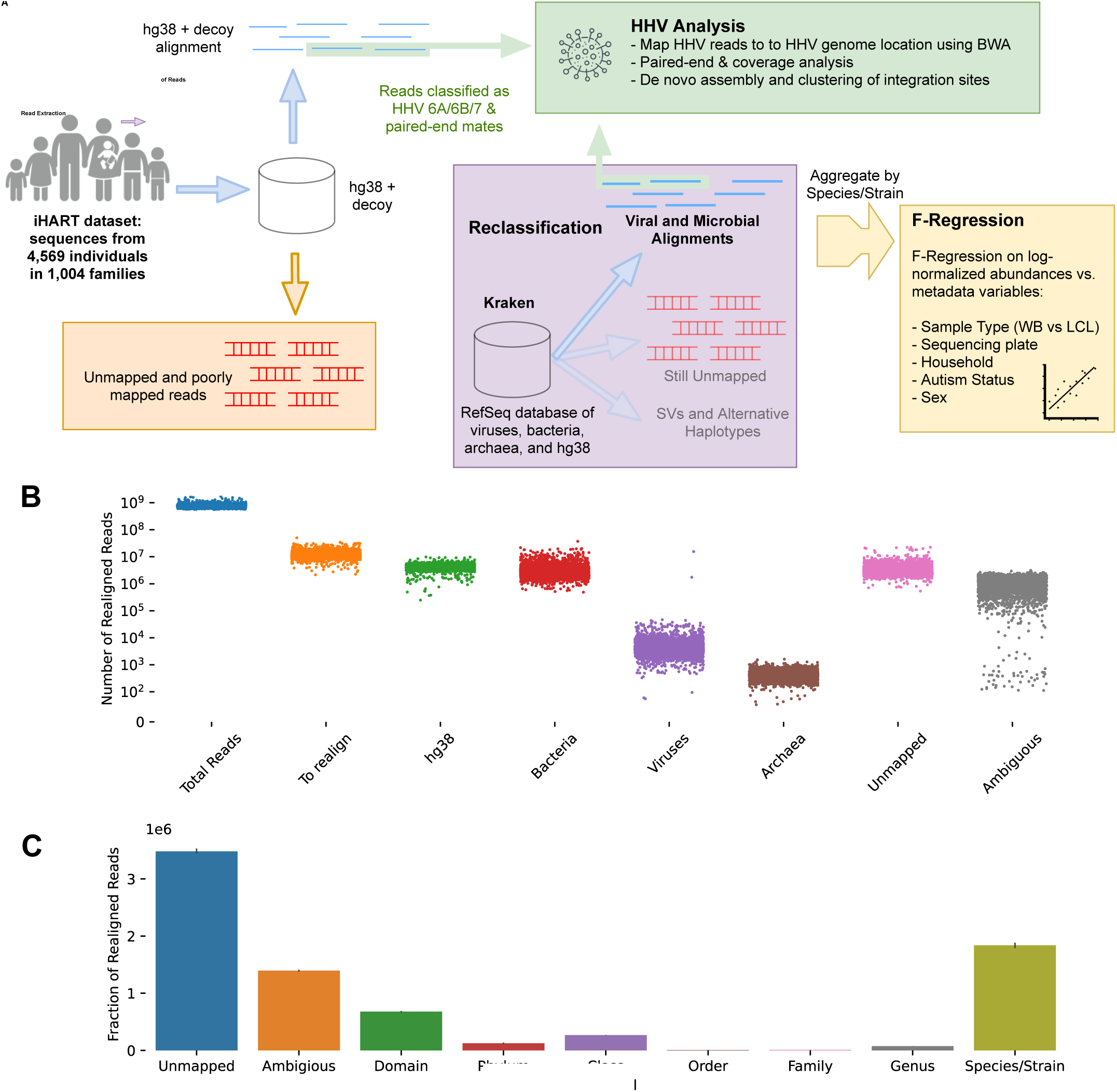
**(A)** General workflow of the study. Poorly mapped and unmapped reads were extracted from the iHART dataset, reclassified to a database of bacteria, archaea, and hg38, and aggregated over species. Reads that were classified as human herpesvirus 6A, 6B, or 7 were mapped to the HHV genomes and analyzed in conjunction with their mates. **(B)** Number of total and poorly mapped or unmapped reads per sample, and the distributions of their reclassification phyla. **(C)** Phyla of Kraken reclassifications. Because Kraken classified the majority of reads down to the species level, we aggregated read counts by species/strain.

**Figure 2:**
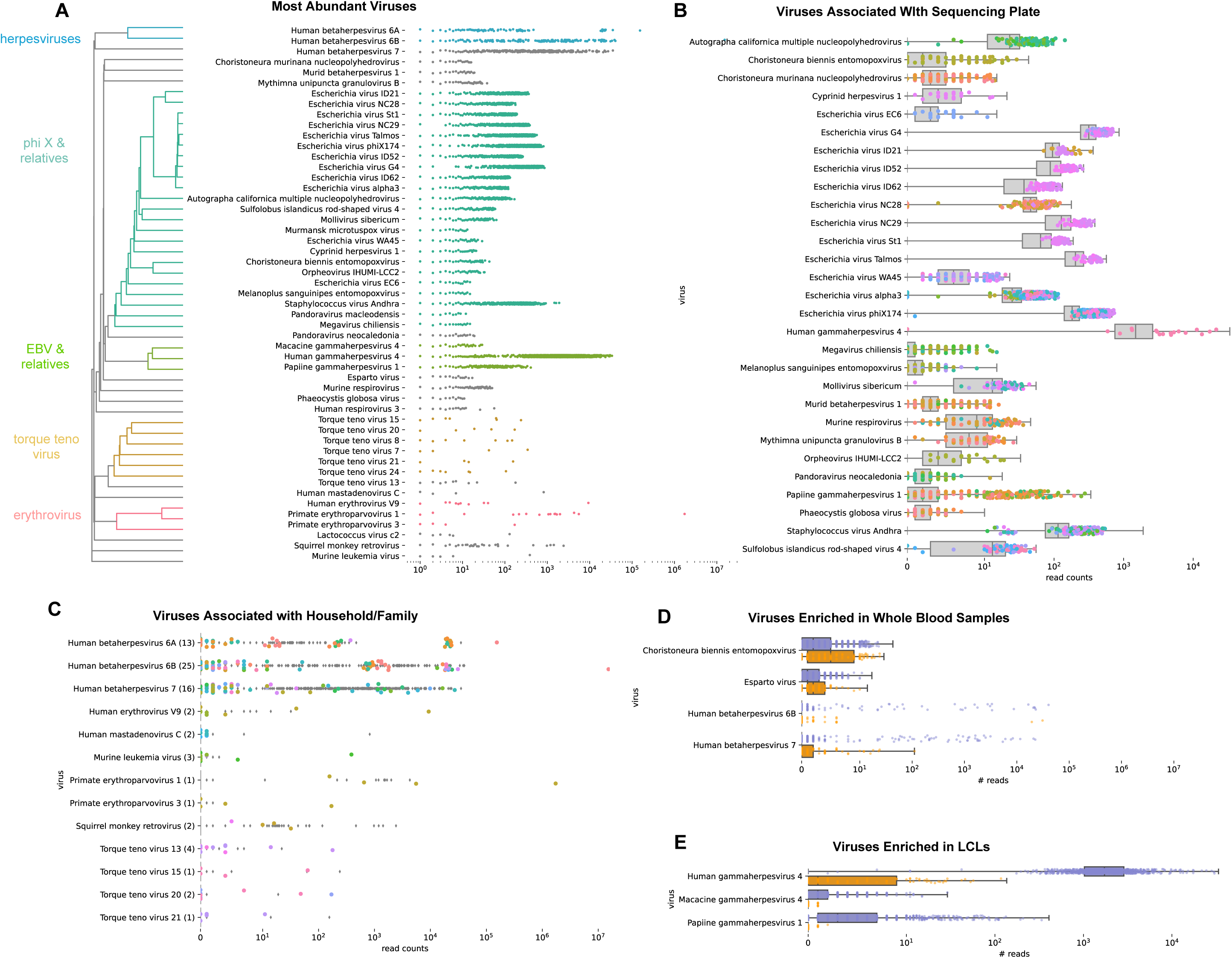
**(A)** Top 50 most abundant viruses, clustered by Spearman correlation across samples. **(B)**Viruses with abundance significantly associated with sequencing plate, per F-regression results. Colors represent sequencing plates that had significantly higher abundances of a given microbe compared to the rest of the population. Samples from sequencing plates without significant enrichment of a microbe are captured in the grey box plots. **(C)** Viruses associated with household per F-regression. Like **(B)**, colors represent households that had significantly higher abundances of a given microbe compared to the rest of the population. Samples from households without significant enrichment of a microbe are captured in the grey box plots. **(D)** Viruses associated with cell type per F-regression results, and more abundant in whole blood samples (orange) than LCL samples (purple). **(E)** Viruses associated with cell type per F-regression results, and more abundant in LCL samples than whole blood samples.

### 3.2 Biological sample source and sequencing plate affect viral reagent and contaminant abundances

Using an F-regression, and regressing viral load against sequencing plate, biological sample source (WB vs LCL), sample metadata (such as autism phenotype, sex, household/family, and parent vs. child status), we identified several viruses significantly associated with sequencing plate (Fig. 2B), biological sample type (Fig. 2D-E), and household/family (Fig. 2C).

One reason viruses may show different levels of abundances between biological sources is because they preferentially infect lymphocytes or a different type of blood cell. This is likely the case for TTV and human herpesviruses, both of which have been shown to preferentially infect specific types of blood cells over others [47, 2]. Another reason for biological source-dependent abundance is that different reagents and pipelines are used in LCL versus whole blood prep and storage pipelines and may lead to different contamination profiles. This is probably the case for EBV (gamma herpes 4) enriched in LCLs (acquired during the EBV-induced immortalization step) and its relatives, as well as the non-herpesviruses enriched in whole blood samples. Similarly, viruses enriched in specific sequencing plates (Fig. 2B) indicate batch contamination introduced during sequencing prep and sequencing.

### 3.3 Several Viruses Show Evidence of Household Transmission

Interestingly, we found several viruses associated with household/family, indicating that family members may be transmitting an active infection within their household. Even in low counts, we see a statistically significant family association for torque teno virus, as well as for erythroparvovirus. We also see an association between family and human herpesviruses. This particular association is likely driven by two mechanisms: inherited chromosomally integrated human herpesvirus (iciHHV), which is passed down from parent to child through Mendelian inheritance, and active infections being transmitted within a household. We discuss the differences in viral load profiles of the different human herpesviruses later on.

### 3.4 0.6% of Population Shows Evidence of iciHHV-6

Human herpesvirus 6 can integrate into host genomes, and ancient germline integration events can be seen in present day as mendellianly inherited genotypes. We identified 28 samples (.6% of samples, 14 with iciHHV-6A and 14 with iciHHV-6B) with a high likelihood of having iciHHV-6A or iciHHV-6B. These samples had HHV read counts consistent with 1 copy of HHV per cell (or .5 HHV genomes/human genome copy), and had a parent or child in the same family also with high HHV-6A or 6B counts. The HHV counts of these samples and others are shown in Fig. 3A. The probable iciHHV-6B samples came from both WB and LCLs. While all 14 cases of iciHHV-6A were only found LCL samples, the LCL samples outnumber WB by 10-fold so this was not statistically significant. There was no overlap between the samples with likely iciHHV-6A and those with likely iciHHV-6B. Additionally, no samples showed evidence of homozygous iciHHV (a copy of iciHHV inherited from each parent).

**Figure 3:**
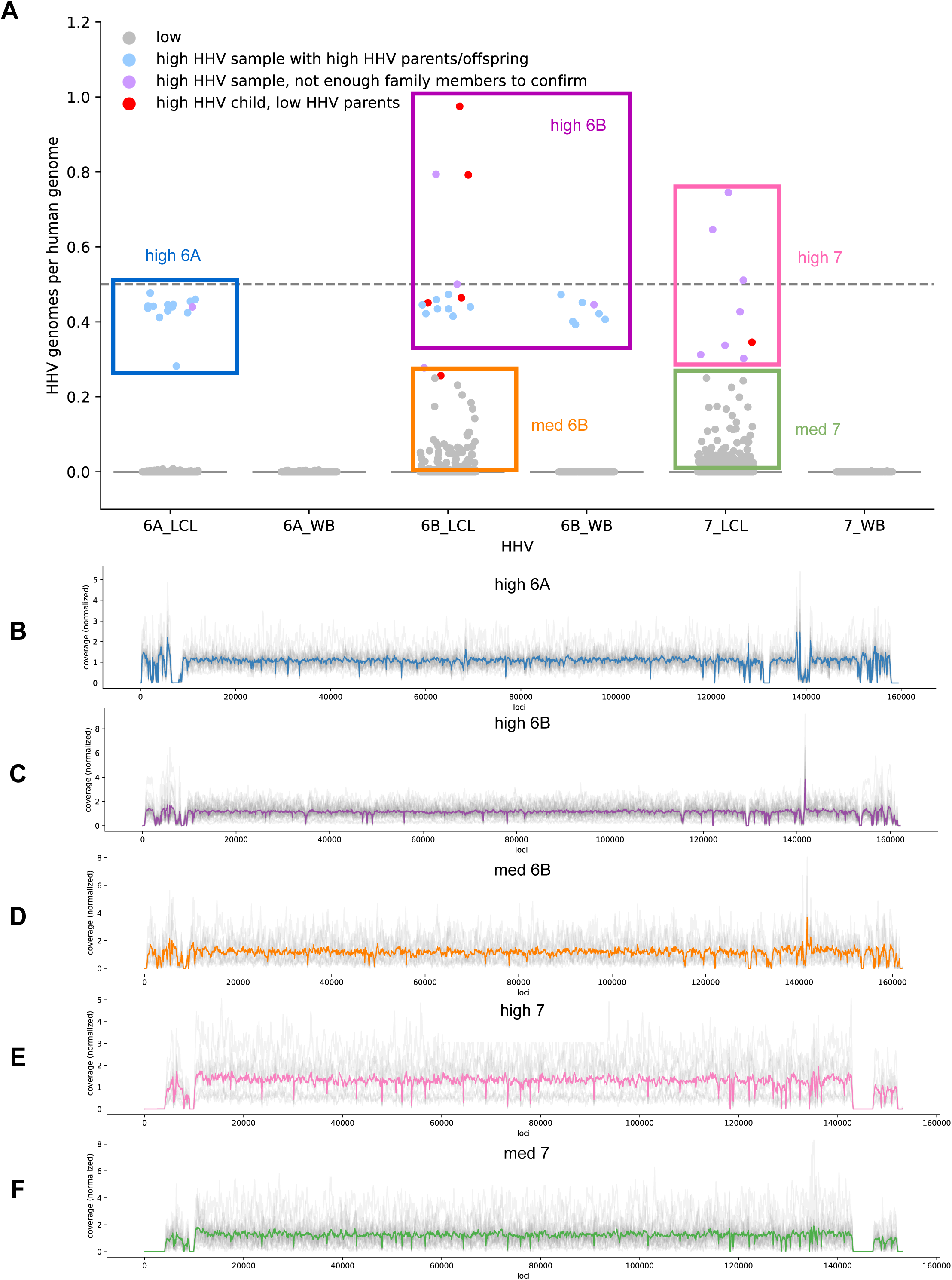
**(A)** Counts of HHV genomes, normalized by coverage of housekeeping genes ERB and HBB. A normalized HHV genome/human genome fraction of around .5 is consistent 1 HHV genome per host cell, and an HHV genome/human genome fraction of 1 is consistent with 2 HHV genomes/host cell. Dots represent samples and are colored by observed inheritance patterns, and several groups of interest are boxed. **(B)-(H)** Normalized coverages of HHV genomes. A normalized coverage of 1 indicates that the expected number of HHV reads under a uniform coverage distribution fall under that region of the genome. Grey lines indicate coverages distributions of each sample, and bold colored lines represent the average. **(B)** Normalized coverage of HHV-6A from samples with “high 6A” defined by > .25 HHV-6A genomes/human genome. **(C)** Normalized coverage of HHV-6A from samples with “high 6B” defined by > .25 HHV-6B genomes/human genome. **(D)** Normalized coverage of HHV-6B from samples with “medium 6B” defined by between .01 and .25 HHV-6B genomes/human genome. **(E)** Normalized coverage of HHV-7 from samples with “high 7” defined by > .25 HHV-7 genomes/human genome. **(F)** Normalized coverage of HHV-7 from samples with “medium 7” defined by between .01 and .25 HHV-7 genomes/human genome.

Additional evidence for iciHHV-6 comes from the coverage profiles of samples with high HHV-6A and high HHV-6B (Fig. 3B-C). Although there are some regions with slightly lower or higher average coverages (corresponding to homologous regions between 6A and 6B, low complexity regions, or high GC content regions), no single region dominates the dominates the coverage profile, indicating that full HHV-6 viral genomes exist in these samples and are not artifacts of mismappings. Similar coverage profiles were found for HHV-6B with medium viral loads (Fig. 3D) and HHV-7 in samples with both high and medium viral loads (Fig. 3 E-F).

### 3.5 Circulating HHV-6B De Novo integrates into lymphocytes

While HHV-6A abundance showed a clear bimodal distribution, HHV-6B had a more continuous distribution in the LCLs, with many samples having abundances ¿.1 copies/genome reads but not showing inheritance patterns consistent with iciHHV. Thus, HHV-6B has a chromosomal integration pattern that suggests two distinct types of integration events: iciHHV-6 and *de novo* integration events in lymphocytes. We note that the coverage profiles suggest only minimal mismappings between HHV-6A and HHV-6B at conserved regions, and no mismappings from or correlation to gammaherpesvirus 4 **??**.

We found that reads mapped to the end of HHV-6A and HHV-6B frequently had a mate mapped to a specific region on the decoy reference genome, chrUn JTFH01000690v1 decoy **??**A-D. This reference sequence is probably an unplaced telomeric sequence, and serves as a HHV integration site. *De novo* assembling and clustering these potential integration sites, we see common integration sequences for HHV-6A and HHV-6B. In HHV-6B, both probable iciHHV and *de novo* integrated samples share 3 canonical integration sequences.

While LCLs are immortalized by injecting Epstein Barr Virus (herpesvirus 4), HHV-6 does not play a role in the LCL pipeline, nor did we find any relationship between HHV-4 and HHV-6B viral loads. It has been shown that HHV-6 infects lymphocytes, and establishes latency via chromosomal integration[17]. HHV-6B is also extremely prevalent in most populations, more so than the much rarer HHV-6A [14]. Therefore, a plausible explanation for this spectrum of HHV-6B abundance is that HHV-6B is chromosomally integrated with a fraction of the cells in the sample: During a HHV-6B infection, HHV-6B established latency via integration one or more lymphocytes. Genetic drift, natural selection, or reactivation during that person’s life and during LCL passaging causes different samples to have different fractions of infected cells.

### 3.6 Circulating HHV-7 reactivates and infects lymphocytes

HHV-7 did not show any evidence of iciHHV: no WB cells had high counts of HHV-7, and none of the LCL samples with HHV-7 counts consistent with iciHHV have parent-offspring relationships. This is consistent with findings that HHV-6, but not HHV-7, can integrate into host chromosomes [26]. However, like HHV-6B, HHV-7 shows a continuous distribution in the LCLs, suggesting that HHV-7 latency combined with genetic drift, positive selection, or reactivation causes different samples to have different fractions of HHV-7 genomes. On the other hand, HHV-7 reads did not consistently pair with any hg38 or decoy contig, nor did they frequently pair with unmapped reads. This is in agreement with the differences between HHV-6 and HHV-7 infection: while HHV-6 establishes latency primarily via chromosomal integration, HHV-7 establishes latency by forming an episome inside the host nucleus [14]. We thus hypothesize that a primary infection causes a small fraction of lymphocytes in sample to initially contain latent HHV-7. Either through replication of the episome, or reactivation and infection of additional cells in the sample, HHV-7 increases its load throughout LCL immortalization and passaging.

Interestingly, we ran the same HHV alignment pipeline on unmapped reads from the 1000genomes dataset [38] of high coverage WGS from around the world. We did not find a continuous distribution of HHV-6B; rather we found a bimodal distribution with most samples having almost 0 HHV-6B read counts, and <1% of samples having HHV-6B read counts consistent with HHV-6B. In the 1000genomes cohort, also WGS derived from LCLs, we also found only one case of medium abundance HHV-6B (500 reads), with the rest of the samples having <10 reads aligning to HHV-7. Notably, we found HHV-7 and HHV-6B to be more abundant in children than parents in our dataset. Because the 1000genomes data we used was all from adults, we hypothesize that childhood infection (coupled with *de novo* integration or latent establihsment of HHV-6B and HHV-7 into lymphocytes) is driving the odd distributions of HHV-6B and HHV-7 in the iHART dataset. Alternatively, the immortalization and storage processes in the iHART dataset may be increasing integration and intra-sample re-infection rates.

### 3.7 HHV Displays Genetic Diversity across Hosts

We wished to understand the origins and diversity of circulating, latent, and iciHHV. We *de novo* reconstructed the HHV genome from each sample where possible (*de novo* assembly failed for samples with low HHV read counts), and compared genomes using MAAFT multiple sequence alignment and ClustalW phylogenetic tree generation (See Methods). As seen in Fig. 4, HHV 6A, 6B and 7 also exhibit genetic diversity across our samples. HHV 6A genomes fall into three distinct clusters. Family members always fall into the same clade, presumably because these are cases of iciHHV, and parents always pass on the same variant of HHV to their offspring. HHV-6B also exhibits genetic diversity, with genomes in many different clusters that are less distinct than those of HHV-6A. Notably, samples with likely iciHHV-6B do always fall into the same clade as their family members, an the HHV genomes from these families are also very closely phylogenetically related to each other. HHV-7 also exhibits genetic diversity, and does not seem to originate from a single source (as might be the case if HHV-7 was a contaminant). Interestingly, HHV-7 genomes from members of the same family tended to be much closer phylogenetically than HHV-7 from unrelated individuals (Mann-Whitney U test using distance matrix values, p-value < .05). Removing the suspected iciHHV cases, HHV-6B also showed the same trend (p-value < .05). This may indicate that the HHV-6B and HHV-7 variant that established itself in LCLs originated from an initial infection that was spread within a household.

**Figure 4:**
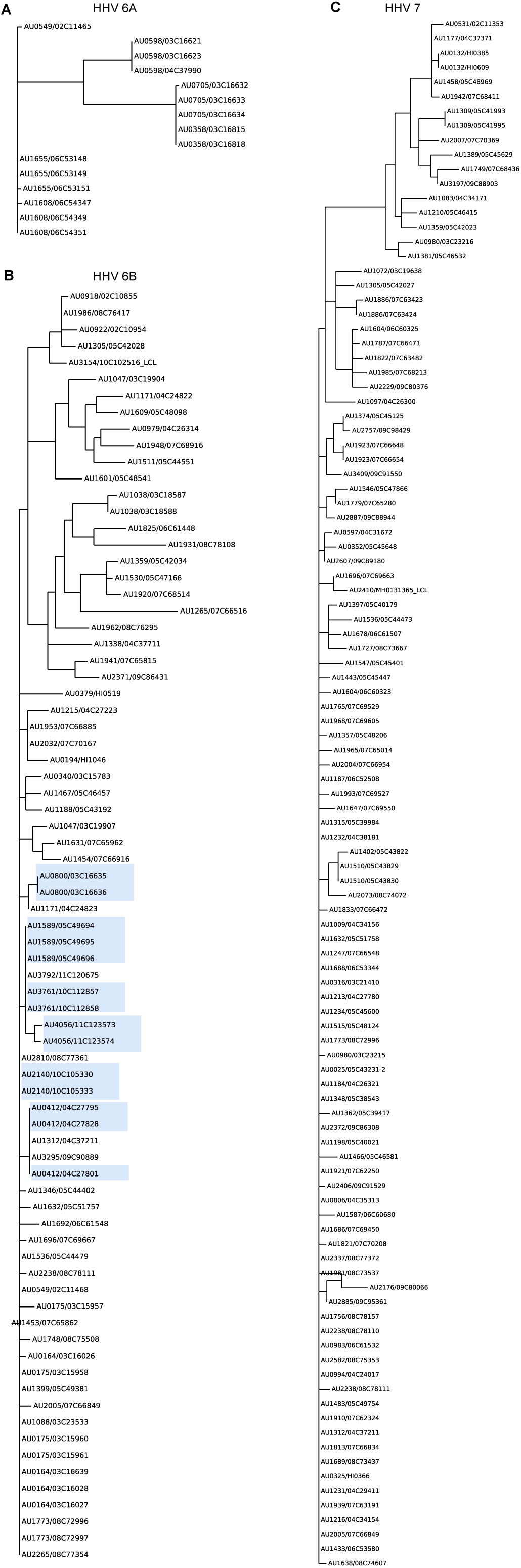
Phylogenetic tree of *de novo* assembled HHV genomes from the samples where *de novo* assembly was successful. Leaves are labelled with family and sample IDs <FAMILY ID>/<SAMPLE ID>. **A** Phylogenetic tree of HHV-6A. **B** Phylogenetic tree of HHV-6B, with suspected cases of HHV-6B highlighted in blue. **C** Phylogenetic tree of HHV-7

### 3.8 Several Canonical Telomeric Sequences are Integration Sites for HHV 6A or 6B

Using megahit, MAAFT, and ClustalW, we *de novo* assembled, aligned, and built a phylogenetic tree from the reads that did not align to HHV-6A, 6B, or 7 but had mates that aligned to HHV. HHV-7 had very few of such reads and thus *de novo* assembly was not possible in any sample. However, HHV-6A and HHV-6B show clear canonical flanking sequences, which we refer to as candidate integration sites (Fig. **??**). Interestingly, there is little variation within the 5’ and 3’ integration sites for HHV-6A. Small single-nucleotide differences are shared among family members, indicating inherited integrated viruses and sites.

HHV-6B 5’ and 3’ flanking regions also cluster into clear canonical candidate integration sites. Both the 3’ and 5’ sites cluster into 3 distinct clusters, with highly dissimilar sequences. Family members with suspected iciHHV-6B usually fall within the same cluster, however in the 5’ flanking integration site families AU0412, AU2140, and AU4056 fall into separate clusters and in the 3’ flanking region members from family AU4056 falls into separate clusters.

When we matched the candidate integration sites to public sequences using NCBI’s BLAST, all sequences matched to isolate HHV or endogenous HHV sequences. In particular, sequences matched to studies studying integrated HHV diversity [3, 56, 57, 21, 23].

## 4 Discussion

### 4.1 Viral Transmission and Diversity Within Families

Using whole genome sequences, we extensively catalogue DNA viruses present in human whole blood and lymphocytes. Additionally, we found several viruses that are often transmitted within families. In particular, erythroviruses and torque teno viruses may be transmitted within households though the mechanism of the particular transmissions in our dataset remains unknown. Previous studies have identified both transplacental and fecal-oral modes of transmission in torque teno viruses [48]. Erythroviruses also can be transmitted transplacentally, and more commonly through respiratory droplets [28]. We additionally identified 28 cases of suspected iciHHV-6. We show that integrated herpesviruses are genetically diverse, with variable genomes sites and integration sites across families. Additionally, herpesviruses seem to be often transmitted within families, as samples from family members more often contain the same exogenous HHV-6B and HHV-7 variant than those from unrelated individuals. It may also be that common variants within families are the result of variation in herpesviruses specific to different regions in the U.S. [21]. HHV has been implicated in several diseases such as multiple sclerosis, encephalomyelitis, and febrile convulsions [13]. Genetic differences in exogenous HHV and iciHHV and its integration sites could influence disease pathology and contribute to different incidences of disease across different regions of the world [49] The unmapped read space of whole genome sequencing data is an easy method for better understanding HHV diversity and its possible role in disease.

### 4.2 Establishment of HHV-6B and 7 into LCLs

To our knowledge, this is the first study to show evidence of widespread replication of non-inherited HHV-6B and HHV-7 in LCLs using thousands of samples. Moreover, previous studies have shown that HHV-6 and HHV-7 typically preferentially infect (CD4+) T-lymphocytes. However, the LCLs from the iHART dataset are derived from B-lymphocytes, indicating that B-lymphocytes may be an underappreciated route for HHV infection.

We hypothesize that the replication of HHV-6B and HHV-7 occur by separate but related mechanisms. We hypothesize a primary infection of HHV-6B in one or more lymphocytes from the donor *de novo* integrated into the host chromosomes (as evidenced by the reconstructed integration sequences), either while still in the host or during the process of LCL immortalization and storage. Genetic drift, positive selection, or reactivation then increased the fraction of cells with an integrated virus over time, leading to varying loads of HHV-6B across samples. HHV-7 on the other hand, is not known to chromosomally integrate, but rather establishes latency via an episome inside the nucleus. Similar to HHV-6B, we hypothesize a primary infection of HHV-7 results in an HHV-7 episome inside a small fraction of cells. The episome replicates in tandem with the host chromosomes, Again, genetic drift of positive selection increases the fraction of cells with episomal HHV-7. This would be very similar to the life cycle of Karposi’s sarcoma herpesvirus (HHV-8) [24], which establishes latency via an extrachromosomal nuclear episome that colocalizes to the chromosomes in order to replicate in tandem with the host cell.

### 4.3 Potential of the Unmapped Read Space for Surveillance of Pathogens

In this study, we have used the unmapped read space of whole genome sequences to better understand prevalence and intra-family transmission patterns of various blood viruses. To our knowledge this is the first study using large WGS datasets of families in order to study viral transmission. Additionally, the unique family structure of our dataset allowed us to identify likely cases of iciHHV-6A and iciHHV-6B and document the genetic and integration site variation within these species. This is also the first study to observe and hypothesize about the widespread *de novo* HHV-6B integration and HHV-7 replication in LCLs. We hope this encourages further research on HHV-6 and HHV-7 integration and latency. The samples in our dataset with these unique distributions are available for future research upon request and application.

We performed such analyses using a collection of WGS data that was generated for unrelated purposes (to understand the genetic components of autism). We suspect whole genome sequences contain a wealth of untapped data, and may be valuable resources beyond their traditional GWAS use cases. Particularly, as more WGS data is generated from diverse global populations, the unmapped read space could be used to track the spread and geography of various viruses.

## 5 Methods

### 5.1 Dataset and Original Alignment to hg38

We obtained Whole Genome Sequencing (WGS) data from the Hartwell Autism Research and Technology Initiative (iHART) database, which includes 4,842 individuals from 1,050 multiplex families in the Autism Genetic Resource Exchange (AGRE) program [44]. A total of 4,568 individuals from 1,004 families passed quality control and were included in the analyses. DNA samples were derived from whole blood (WB) or lymphoblastoid cell lines (LCL) and sequenced at the New York Genome Center.

All WGS data from the iHART database have been previously processed using a standard bioinformatics pipeline which follows GATK’s best practices workflows. Raw reads were aligned to the human reference genome build 38 (GRCh38 full analysis set plus decoy hla.fa) using Burrows-Wheeler Aligner (bwa-mem).

### 5.2 Extracting Unmapped and Poorly Unmapped Reads

We excluded secondary alignments, supplementary alignments, and PCR duplicates from downstream analyses. We extracted reads from the iHART genomes that were unmapped to hg38 and the decoy reference or mapped with low confidence. Low-confidence reads were defined as reads marked as improperly paired and had an alignment score below 100. We used alignment score rather than mapping quality in order to select for reads were likely not true alignments to the human reference genome, rather than for reads that had ambiguous alignments to hg38. These reads were then re-paired if both ends needed to be realigned, and lastly separated into single-end and pair-end files.

### 5.3 Taxonomic Classification and Aggregation

We used Kraken2 [55] to align the unmapped and poorly aligned reads to a the Kraken default (RefSeq) databases of archaeal, bacterial, human (GRCh38.p13), and viral sequences [39]. These references databases were accessed on Feb 16, 2021. Kraken2 was run on the unmapped and poorly mapped reads from each sample, using the default parameters. Because Kraken was able to map the majority of reads down to the species or strain level, Kraken classifications were aggregated by species before downstream analysis.

### 5.4 F-Regression on Metadata

To analyze the effect of various demographic (such as household, autism status, and sex) and experimental parameters (such as sequencing plate and sample type) on microbial and viral profile, we performed an F-regression analysis. We chose an F-regression because many variables were highly collinear with each other: for example, samples from the same household were nearly always sequenced on the same sequencing plate, autism is much more prevalent in males, and the same sample types were normally collected from households. For each microbe, we built an ordinary least squares (OLS) model, using as our regressor an indicator matrix of sample type, sex, child vs. parent, autism status, sequencing plate, household/family, and sample id, and as our response variable the log-normalized counts of microbes (with pseudo-counts of 1). Using the `statsmodels` library, we then ran a forward OLS regression in which we iteratively selected the regressor features that best explained the previous models residuals, and ceased adding features when the ANOVA score between the previous and new models was no longer statistically significant (adjusted p-value<.05).

### 5.5 Realignment to Herpesvirus Reference Genomes

Using bwa-mem with the default parameters, we aligned all reads classified by Kraken as belonging to herpesviruses to a set of reference genomes consisting of hg38 and the decoy, and all the herpesvirus genomes present in the RefSeq database. Most importantly, this included human betaherpesvirus 6A (NC 001664.4), human betaherpesvirus 6B (NC 000898.1), human betaherpesvirus 7 (NC 001716.2), and both the decoy and RefSeq genome for human gammaherpesvirus 4, or the Epstein-Barr virus (chrEBV in the decoy genome, and NC 007605.1 in RefSeq). We performed the same analysis using 2504 high-coverage WGS LCL samples from the most recent release of the 1000 genomes dataset (Fig. **??**).

### 5.6 HHV Read Counts, Paired-end Analysis, and Coverages

To covert the herpesvirus read counts to viral genomes per host genome, we normalized against the average coverage for two housekeeping genes are not known to show copy number variation, EDAR and HBB [43].

We used pysam [1] and an in-house script to collect genome-wide coverages for different combinations of pairings in order to generate the coverage graphs in Fig. 3 and Fig. 5.

**Figure 5:**
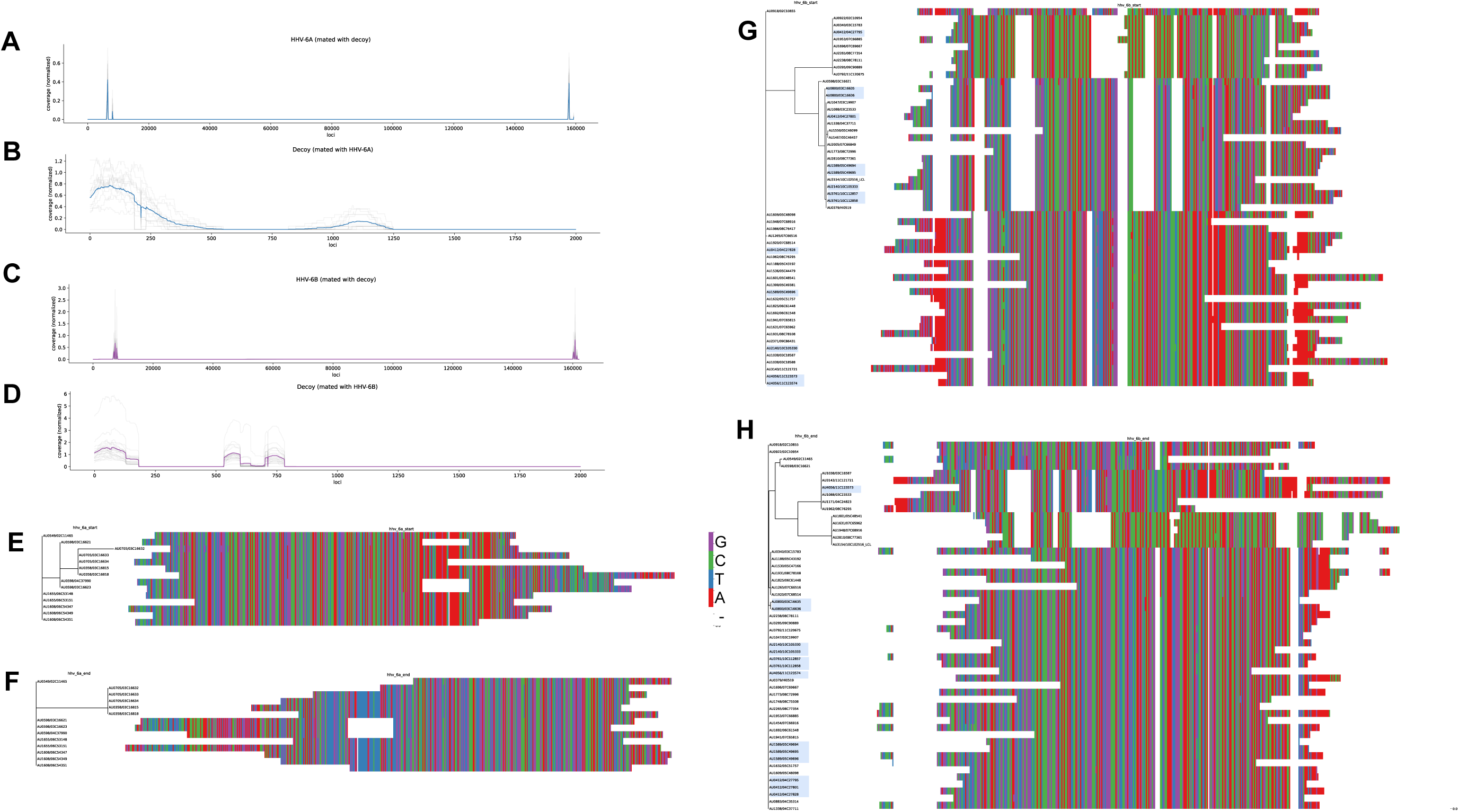
Integration sites and ends of HHV-6A and HHV-6B. **(A)-(D)** Paired coverage analysis. Grey lines indicate the normalized coverages of each sample, and the bold colored line indicates the average. **(A)** and **(B)** were created using samples with at least .25 HHV-6A genomes/host housekeeping genes, and **(C)** and **(D)** were created using samples with at least .25 HHV-6B genomes/host housekeeping genes. **(A)** Normalized coverage of HHV-6A reads that had a mate paired to the decoy reference sequence `chrUn_JTFH01000690v1_decoy`. A normalized coverage of 1 would indicate that 100% of expected number of reads from a given region of the genome had a mate mapped to the decoy genome. **(B)** Normalized coverage of reads mapped to the decoy reference `chrUn_JTFH01000690v1_decoy` with a mate mapped to HHV-6A. sequence. **(C)** Normalized coverage of HHV-6A reads that had a mate paired to the decoy reference sequence `chrUn_JTFH01000690v1_decoy` **(D)** Normalized coverage of HHV-6A reads that had a mate paired to the decoy reference sequence `chrUn_JTFH01000690v1_decoy`. **(E)-(G)** Phylogenetic trees and aligned sequences of assembled integration sequences. **(E)** Assembled sequence of the HHV-6A 5’ integration site. **(F)** Assembled sequence of the HHV-6A 3’ integration site. **(G)** Assembled sequence of the HHV-6B 5’ integration site, with likely cases of iciHHV-6B highlighted in blue.. **(H)** Assembled sequence of the HHV-6B 3’ integration site, with likely cases of iciHHV-6B highlighted in blue.

### 5.7 De Novo Assembly and Clustering of HHV Viral Genomes and Integration Sites

To generate the integration site assemblies and alignments (Fig. 5), we first extracted reads that were not classified as herpesvirus reads but had a mate that aligned to the start or end of the herpesvirus genome. For each individual, we *de novo* assembled these reads. Using MAAFT [**?**], we then performed multiple sequence alignment of these assemblies, and used ClustalW to generate phylogenetic trees. We used the default parameters for MAAFT, and allowed for reverse complementary sequences to be generated as needed. Before generating phylogenetic trees, we attempted to remove redundant sequences that might correspond to a forward sequence and its reverse complementary sequence. We did the following: if a sample had two assembled sequences (presumably corresponding to a forward sequences and a reverse complementary sequence), we removed the sequence that had the least number of matches to the consensus sequence generated by all samples. We used ClustalW on the EMBL browser [30, 34], with a neighbor-joining algorithm, no distance correction, and ignoring gaps.

We BLASTED these assembled sequences against NCBI’s nt nucleotide collection using the default parameters, and not masking low-complexity regions

To generate the assemblies of the viral genomes, we extracted reads aligned to HHV-6A, HHV-6B, and HHV-7. We used `bcftools` to perform variant calling on all of the samples against the reference HHV-6A, HHV-6B, and HHV-7 genomes. We used `VCF2phylip` [41] to convert the variant calls to alternate reference sequences. We filtered to samples that had variants or reference alleles called at at least 50% of loci. Similar to the integration sites, we performed multiple sequence alignment on the reconstructed viral genomes using MAAFT with the default parameters and generated phylogenetic trees using ClustalW using the same parameters as above.

We used Biopython’s Phylo library [9] an in-house python script to generate the sequence alignment trees and diagrams used in Fig. 5 and 4.

### 5.8 Code and Data Access

Analysis code and scripts, as well as the sequences used for KRAKEN classification and herpesvirus realignment can be found at ] https://github.com/briannachrisman/blood_microbiome.

Access to reads from the iHART dataset can be requested via www.ihart.org.

## 6 Acknowledgements

Thank you to The Hartwell Foundation for supporting the creation of the iHART database and the Simons Foundation for additional support for genome sequencing. We thank the New York Genome Center for conducting sequencing and initial quality control of the iHART dataset. We thank Amazon Web Services for their grant support for the computational infrastructure and storage for the iHART database. This work has been supported by grants from The Hartwell Foundation and the NIH (U24 MH081810, R01MH064547, NS101158, NS070911, NS101665, NS095824, S10OD011939, P30AG10161, R01AG17917, and U01AG61356) and from the Stanford Precision Health and Integrated Diagnostics Center and from the Stanford Bio-X Center. Thank you to Jesse Arbuckle and Louis Flamand for their advice and discussions on the HHV distributions.

---

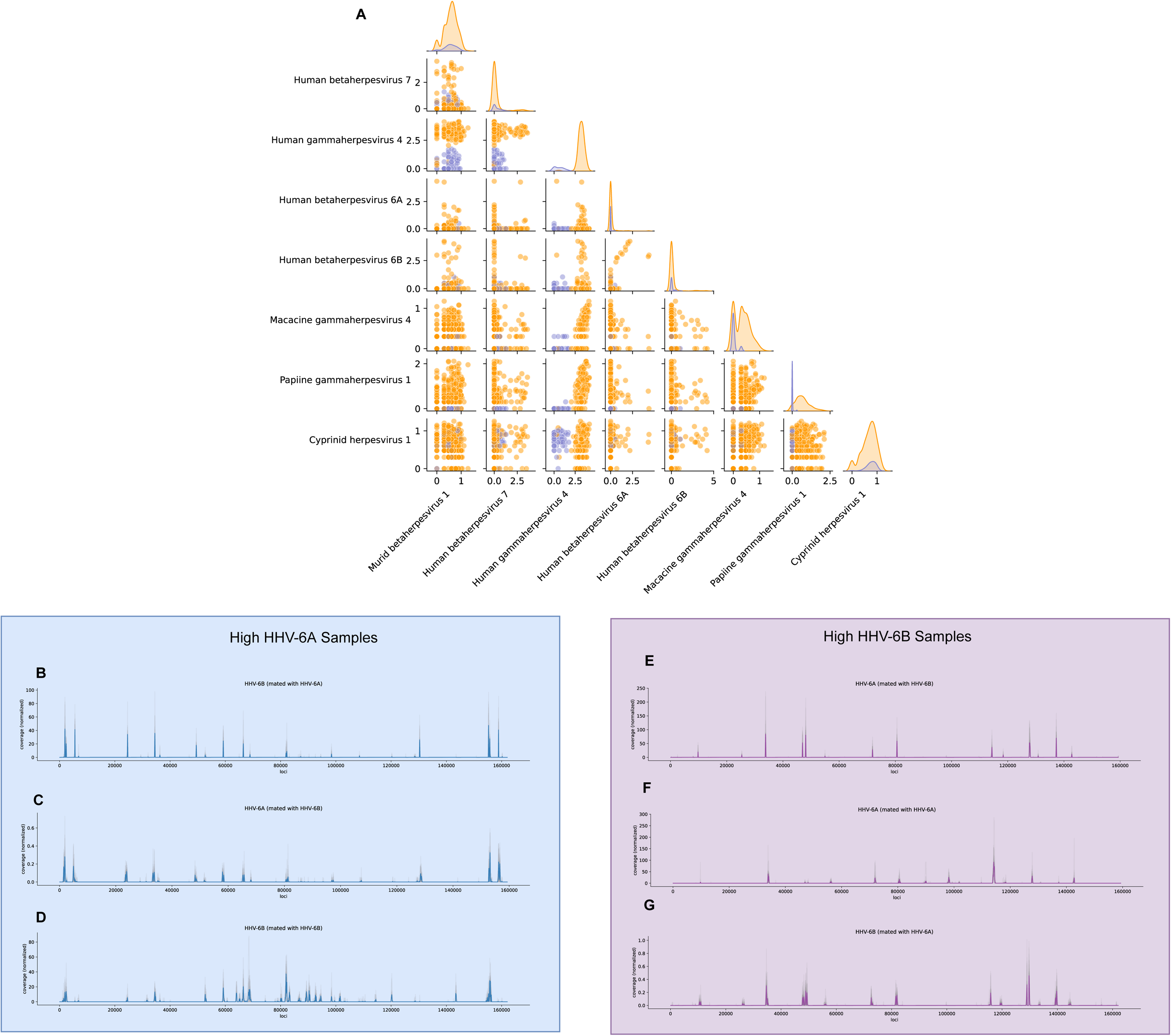

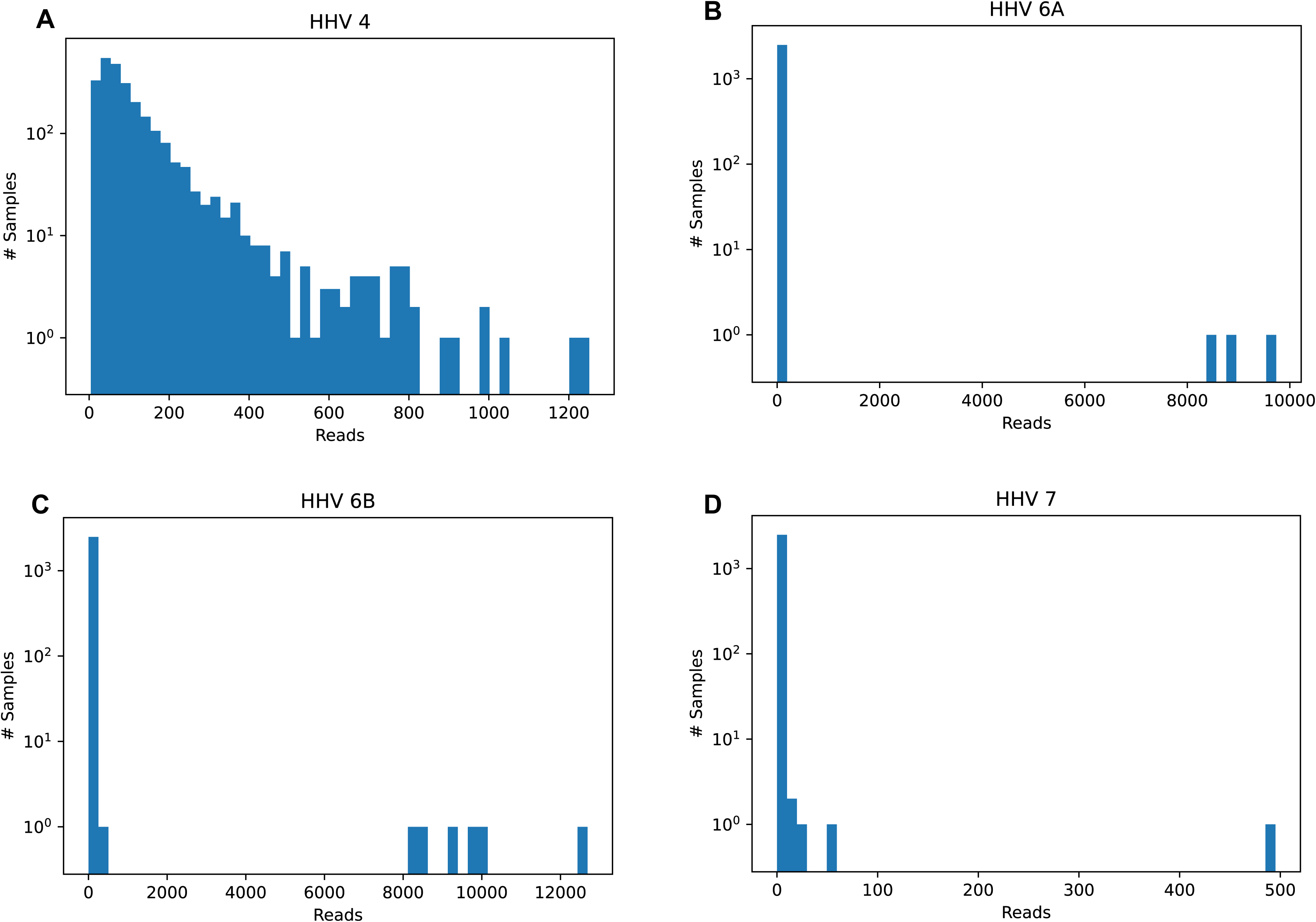

