## Supplementary Figs S1 and S2 for "Transmission Dynamics of Human Herpesviruses and Other Blood DNA Viruses from Whole Genome Sequences of Families"

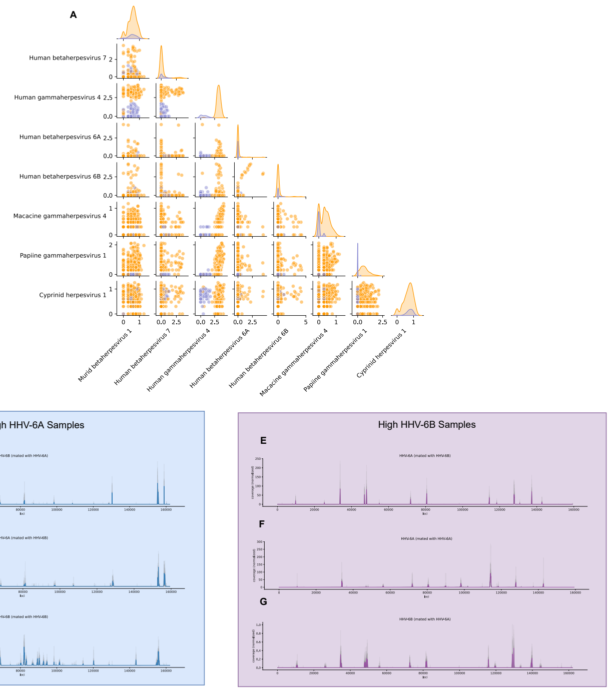


Figure S1: **(A)** Pair-plots of herpesviruses counts. There are some slight correlations due to mismappings between homologous and low-complexity regions. **(B)-(D)** Normalized coverages for samples with high HHV-6A loads. **(B)** Coverage of reads mapped to HHV-6A that were paired with mates mapped to HHV-6B. **(C)** Coverage of reads mapped to HHV-6B that were paired with mates mapped to HHV-6A. **(D)** Coverage of reads mapped to HHV-6A that were paired with mates mapped to HHV-6A. **(E)-(G)** Normalized coverages for samples with high HHV-6B loads. **(E)** Coverage of reads mapped to HHV-6B that were paired with mates mapped to HHV-6A. **(F)** Coverage of reads mapped to HHV-6A that were paired with mates mapped to HHV-6B. **(G)** Coverage of reads mapped to HHV-6A that were paired with mates mapped to HHV-6B.

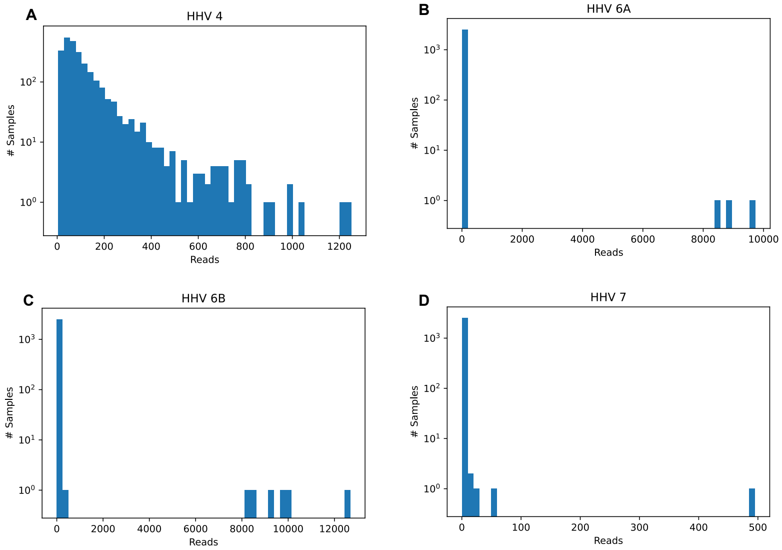


Figure S2: **(A)** Numbers of reads aligned to HHV-4 (Epstein Barr Virus) in the 1000genomes dataset. **(B)** Numbers of reads aligned to HHV-6A in the 1000genomes dataset. **(C)** Numbers of reads aligned to HHV-6B in the 1000genomes dataset. **(D)** Numbers of reads aligned to HHV-7 in the 1000genomes dataset.
